## Supplemental Figures for "Non-coding autoimmune risk variant accelerates T peripheral helper cell development via ICOS"

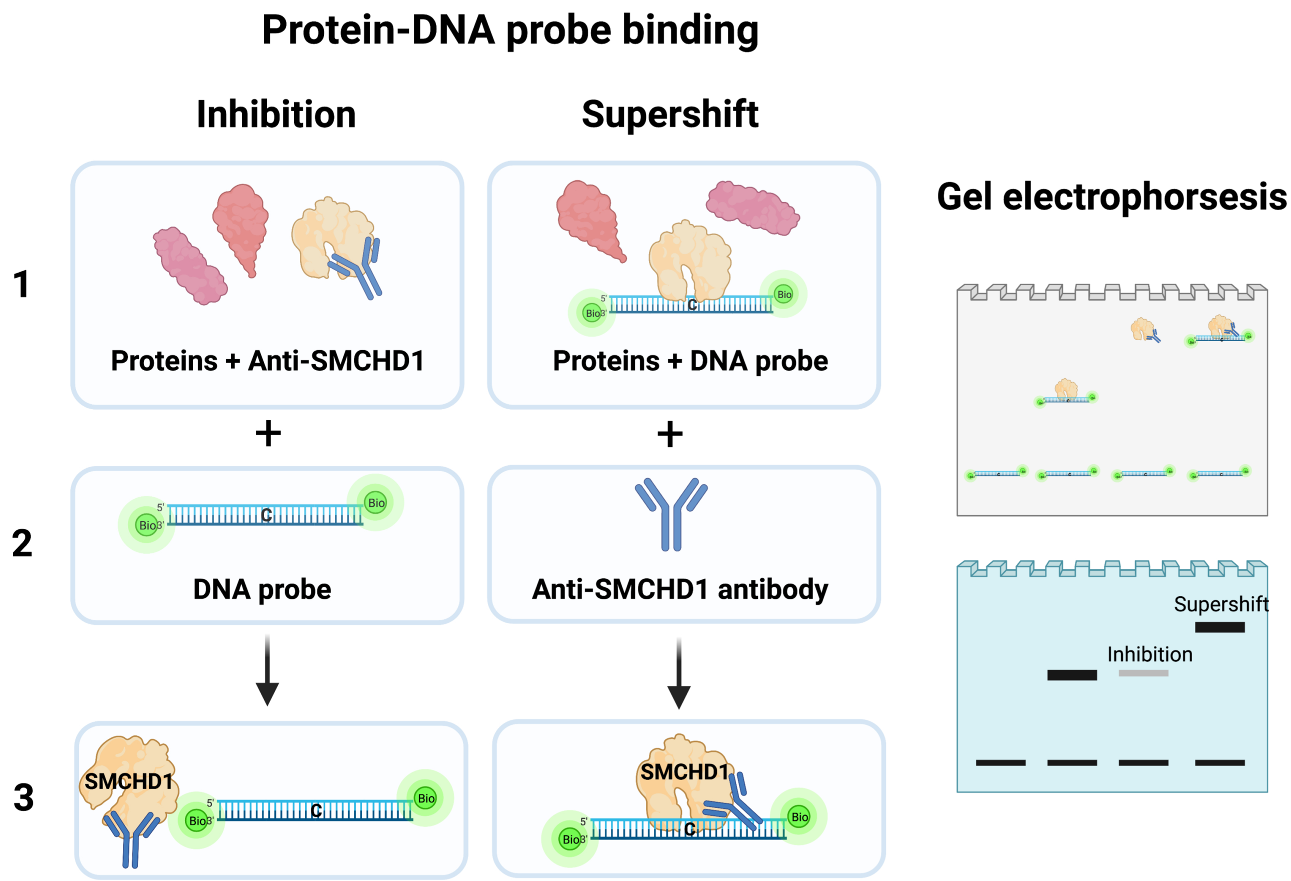


**Supplementary Figure 1**. Differential effects of antibodies during EMSA in Figure 1B. Pre-incubation of anti-SMCHD1 with nuclear extract inhibits DNA probe binding, reducing the amount of SMCHD1-DNA complex observed on the EMSA gel image. Incubation with anti-SMCHD1 after nuclear extract/DNA probe complexes have already been formed results in supershifted complexes of SMCHD1, anti-SMCHD1, and DNA probe.

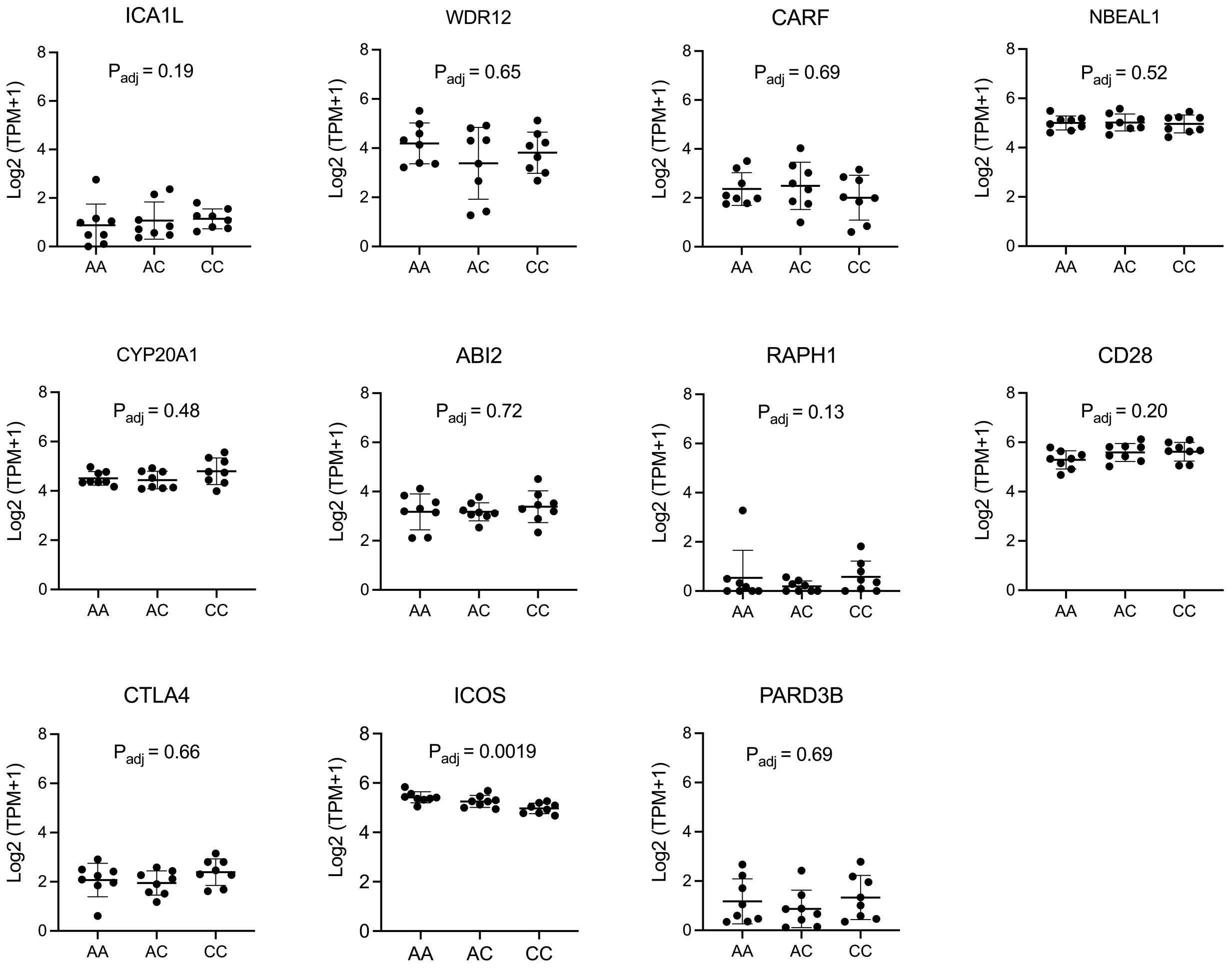


**Supplementary Figure 2**. cis-eQTL mapping analysis was performed using low-input RNA sequencing in resting CD4+ T cells from 24 healthy donors (8 A/A, 8 A/C, and 8 C/C genotypes at rs117701653). We examined 11 protein coding genes that have a transcription start site (TSS) within a 1 MB window of SNP rs117701653, namely *ICA1L*, *WDR12*, *CARF*, *NBEAL1*, *CYP20A1*, *ABI2*, *RAPH1*, *CD28*, *CTLA4*, *ICOS*, and *PARD3B*. Expression levels were corrected for age and sex and rank-normal transformed the residuals. P values from a linear model implemented in QTLtools were shown.

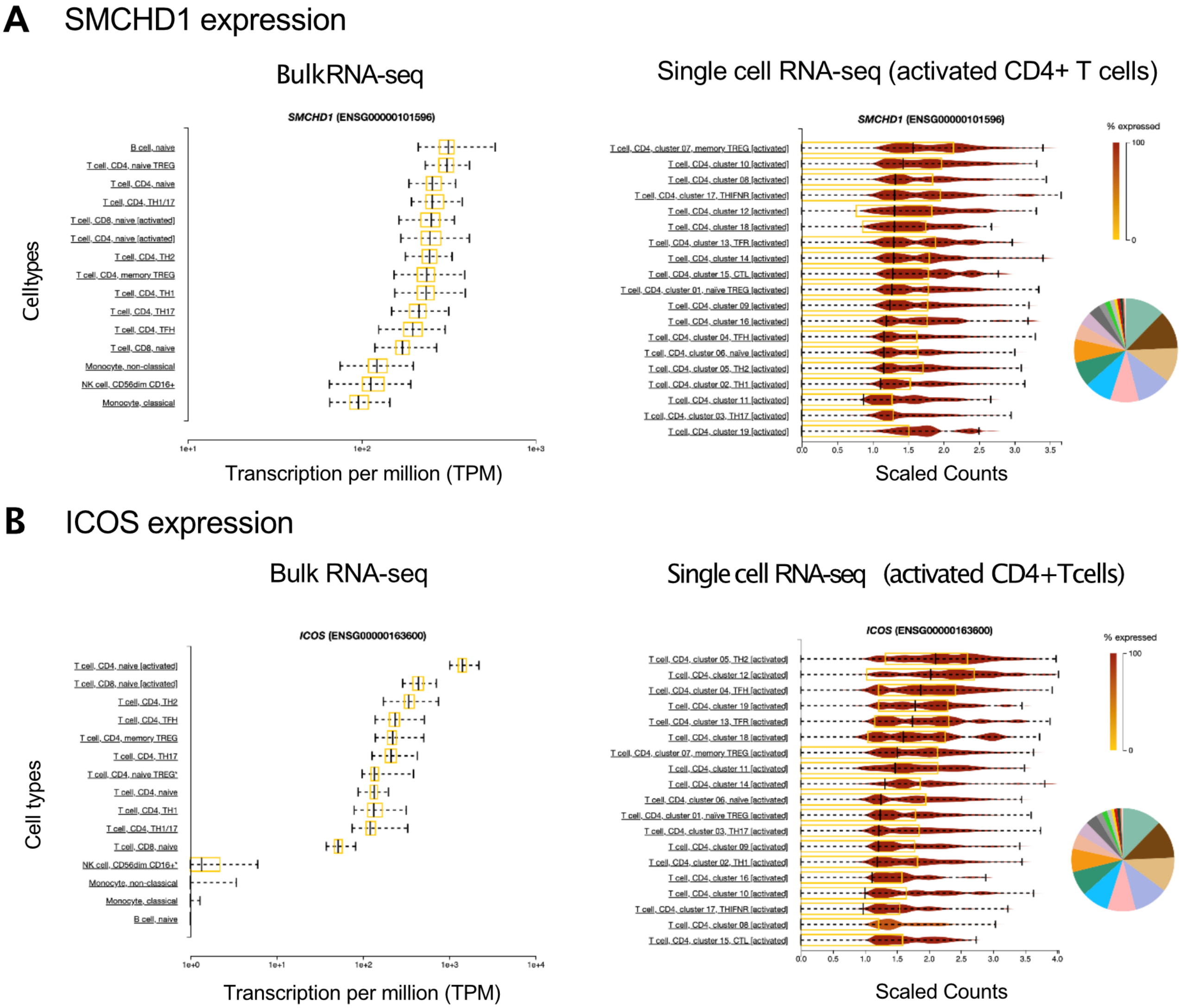


**Supplementary Figure 3.** (A) SMCHD1 and (B) ICOS expressions by bulk RNA sequencing and single cell RNA sequencing were retrieved from https://dice-database.org/landing. Bulk RNA-seq; X-axis represents expression level of depicted genes in transcripts per million (TPM). Boxes indicate 25-75 % interquartile ranges, and whiskers indicate minimum to maximum. Single cell RNA-seq; violin plots display scaled counts of expression levels in each cell cluster. The color scale indicates the fraction of cells that express the depicted genes within each cluster of activated CD4+ T cells. Boxes indicate 25-75 % interquartile ranges, and whiskers indicate minimum to maximum.

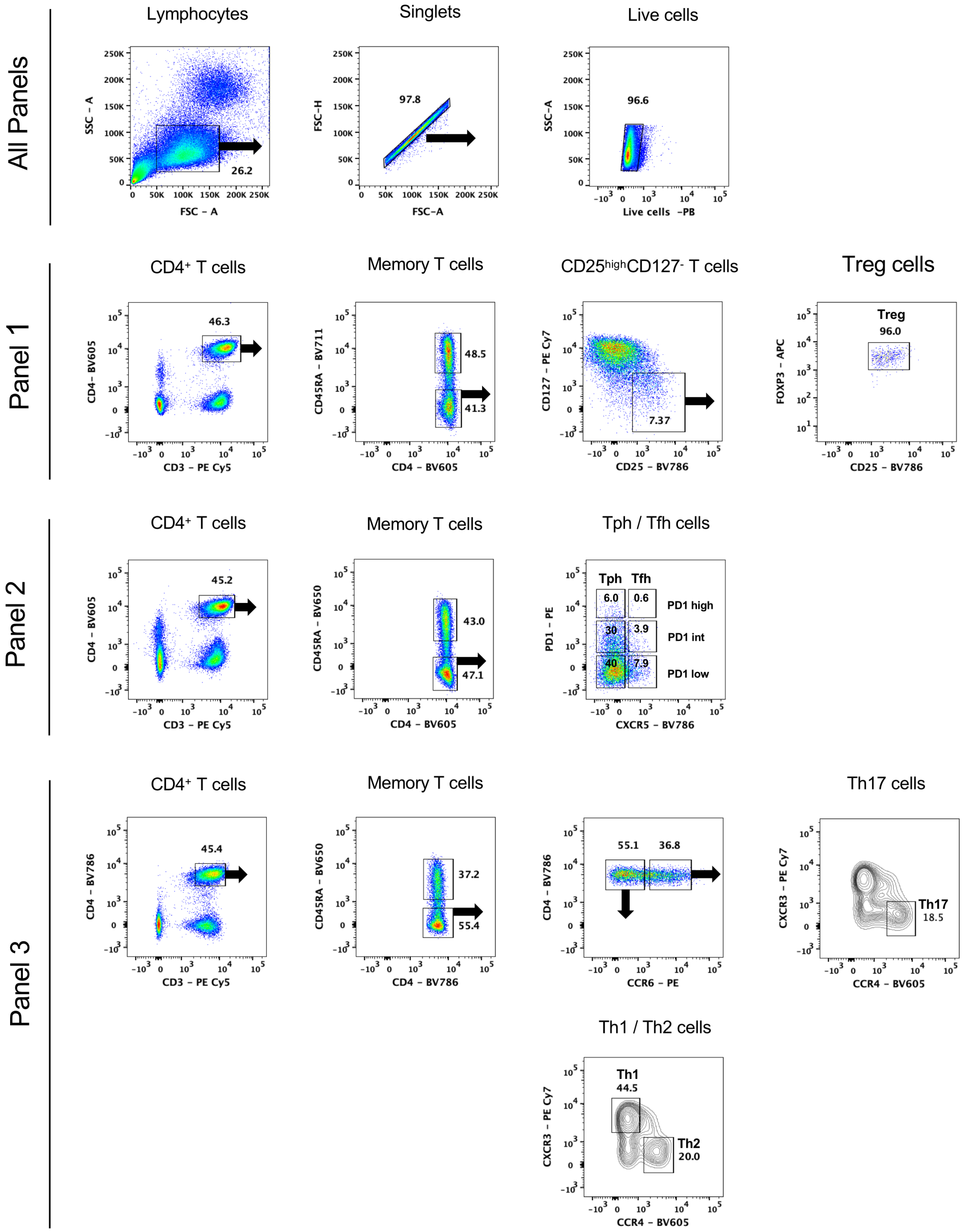


**Supplementary Figure 4.** Gating strategies to identify CD3$^+$CD4$^+$CD25$^{high}$CD127$^-$FOXP3$^+$ Treg cells (panel 1); CD3$^+$CD4$^+$CD45RA$^-$CXCR5$^-$PD-1$^{high}$ Tph cells and CD3$^+$CD4$^+$CD45RA$^-$CXCR5$^+$PD-1$^{high}$ Tfh cells (panel 2); CD3$^+$CD4$^+$CD45RA$^-$ CCR6$^-$CXCR3$^+$CCR4$^-$ Th1 cells, CD3$^+$CD4$^+$CD45RA$^-$CCR6$^-$CXCR3$^-$CCR4$^+$ Th2 cells, and CD3$^+$CD4$^+$CD45RA$^-$CCR6$^+$CXCR3$^-$CCR4$^+$ Th17 cells (panel 3).

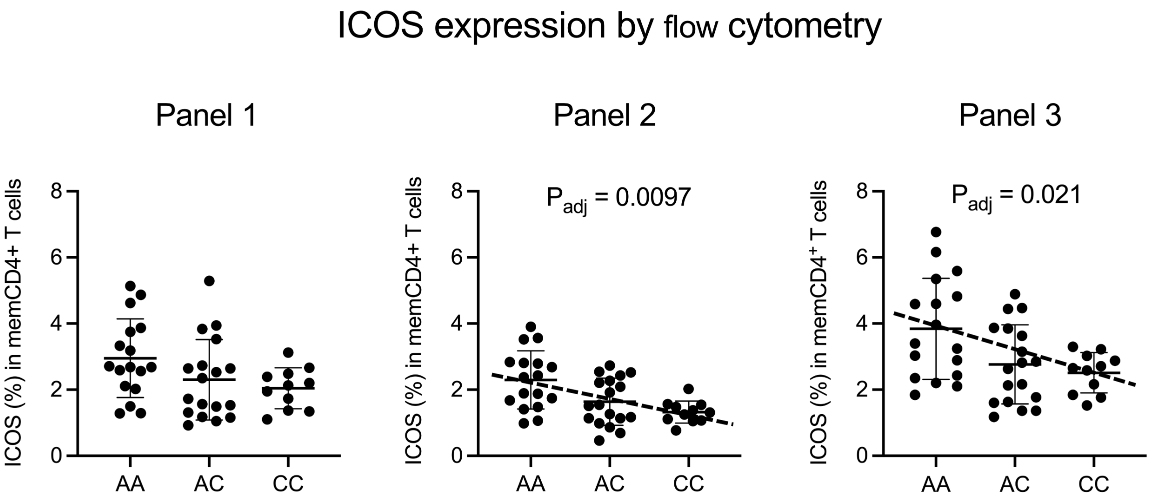


**Supplementary Figure 5**. Association of rs117701653 genotype with frequency of ICOS$^+$ cells among memory CD4$^+$ T cells by flow cytometry analysis, as per antibody panels depicted in Figure S4. Error bars are represented by Mean ± S.D. *P* values were determi ned using a multiple linear regression model adjusted for age and sex.

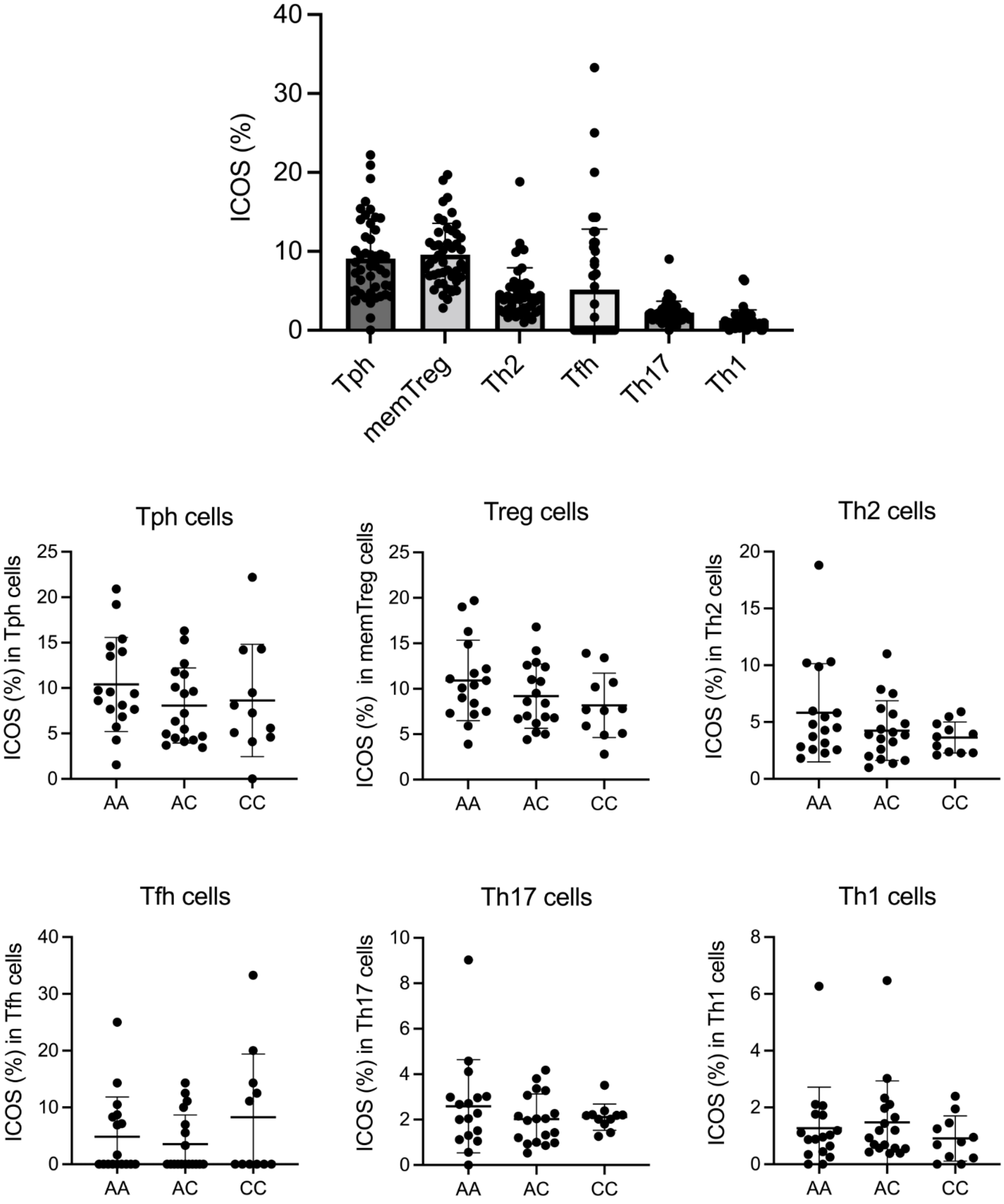


**Supplementary Figure 6**. Frequency of CD4⁺ T cell subsets (Tph, memory Treg, Th2, Tfh, Th17, Th1) that express ICOS was determined by flow cytometry for healthy subjects bearing A/A (n = 17), A/C (n = 18), and C/C (n = 11) genotype at SNP rs117701653.

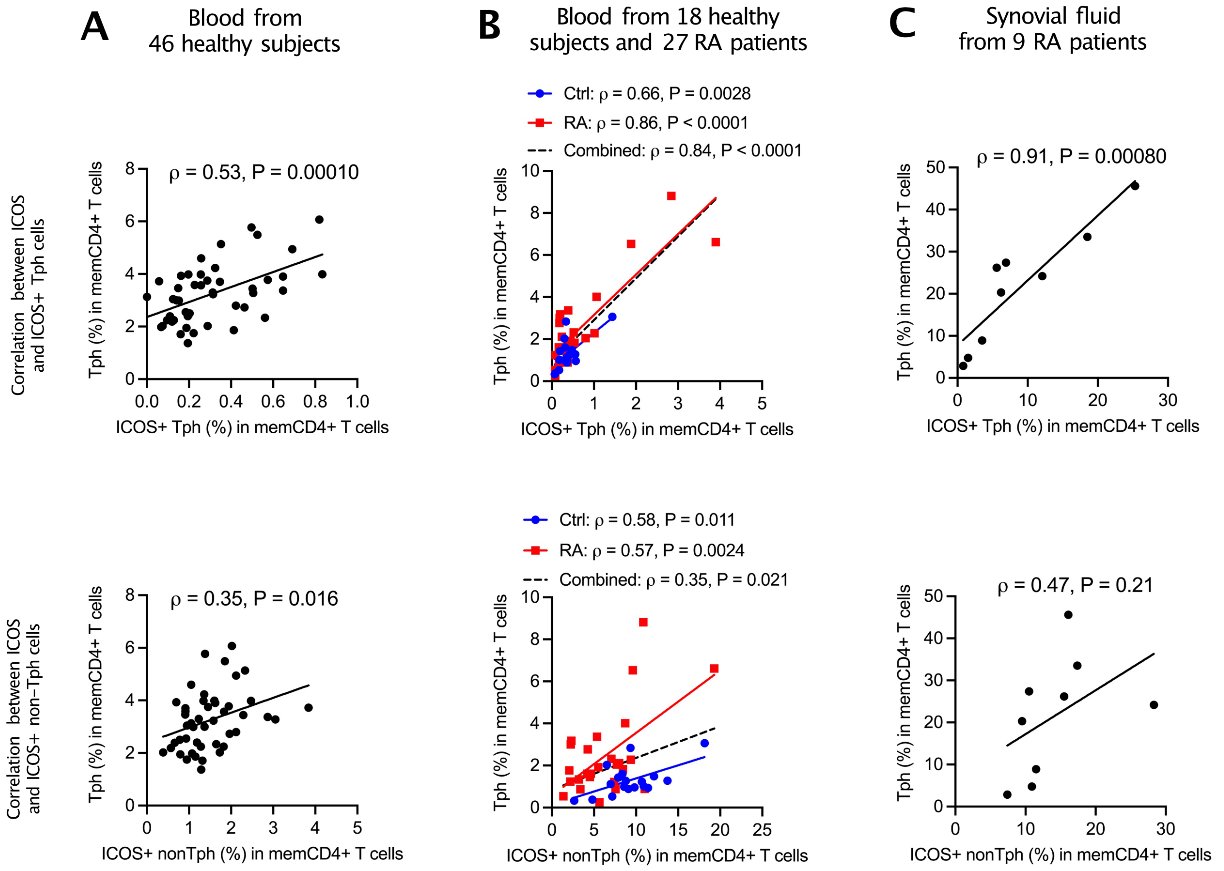


**Supplementary Figure 7**. Frequency of ICOS⁺ Tph or ICOS⁺ non-Tph cells in memory CD4⁺ T cells correlates with frequency of Tph cells in memory CD4⁺ T cells. The proportion of subsets depicted was determined by (**A**) flow cytometry with PBMCs from 46 healthy subjects, (**B**) mass cytometry with PBMCs from 18 healthy controls (colored in blue) and RA patients (colored in red), and (**C**) flow cytometry with synovial fluid from 9 RA patients. Pearson correlation coefficients and *p* values were used to measure linear c orrelations.

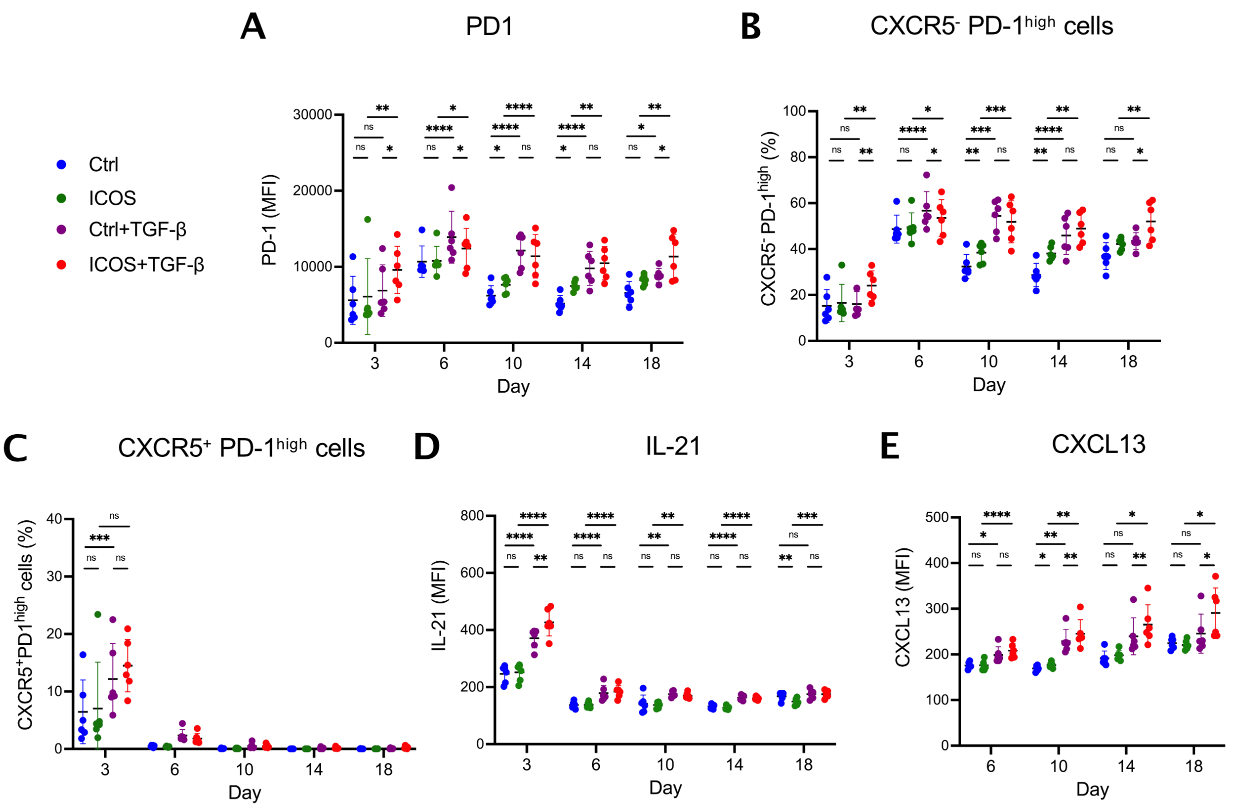


**Supplementary Figure 8.** ICOS stimulation accelerates the development of TGF-β-induc ed CXCR5$^{-}$PD-1$^{high}$ Tph-like cells expressing IL-21 and CXCL13. Human memory CD4$^{+}$ T cells from 6 healthy subjects were differentiated with anti-CD3/CD28 stimulation in the ind icated combination of TGF-β and anti-ICOS stimulation for varying period. Control (Blue), ICOS (Green), TGF-β (Purple), TGF-β+ICOS (Red) stimulation groups were evaluated in time course. (**A**) MFI of PD-1 in whole population. (**B**) Frequency of CXCR5$^{-}$ PD-1$^{high}$ cell s. (**C**) Frequency of CXCR5$^{+}$ PD-1$^{high}$ cells. MFI of (**D**) IL-21 and (**E**) CXCL13 in whole po pulation. MFI, mean fluorescence intensity. Error bars are expressed as mean $\pm$ S.D. *$P < 0.05$, **$P < 0.01$, ***$P < 0.001$, ****$P < 0.0001$. $P$ values were determined using one-wa y ANOVA corrected for multiple comparison by FDR using two-stage linear step-up proce dure of Benjamini, Krieger and Yekutieli.

After 18 days of differentiation

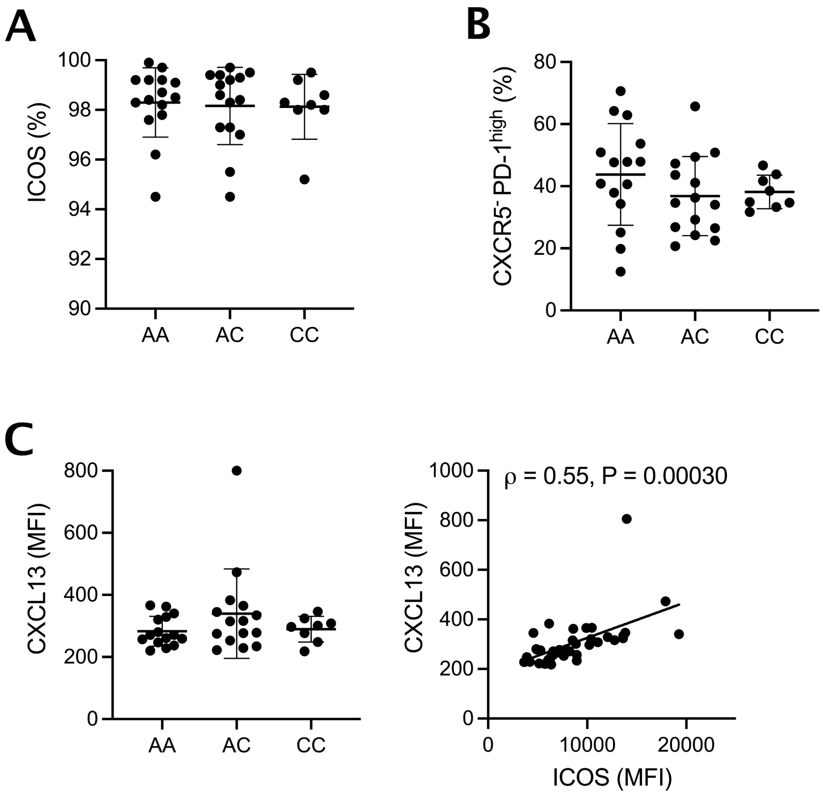


**Supplementary Figure 9.** Genotype at rs117701653 does not correlate with differentiation of CXCR5$^-$PD-1$^{high}$ Tph-like CD4$^+$ T cells at late time (day 18). Memory CD4$^+$ T cells from 38 healthy donors with A/A (n = 15), A/C (n = 15), C/C (n = 8) genotype at rs117701653 SNP were differentiated using anti-CD3/CD28 bead and anti-ICOS antibody in the presence of TGF-β. After 18 days of differentiation, (**A**) frequency of ICOS$^+$ cells, (**B**) frequency of CXCR5$^-$PD-1$^{high}$ cells, (**C**) MFI of CXCL13 and correlation between MFI of ICOS and CXCL13 across whole individuals were displayed. MFI, mean fluorescence intensity. Error bars are expressed as mean ± S.D. *P* values were tested using a multiple linear regression model adjusted for age and sex (**A-C**). Pearson correlation statistics (**C**).
